# Differential gene expression and mitonuclear incompatibilities in fast- and slow-developing inter-population *Tigriopus californicus* hybrids

**DOI:** 10.1101/2022.09.09.507197

**Authors:** Timothy M. Healy, Ronald S. Burton

## Abstract

Mitochondrial functions are intimately reliant on proteins and RNAs encoded in both the nuclear and mitochondrial genomes, leading to inter-genomic coevolution within taxa. Hybridization can break apart coevolved mitonuclear genotypes, resulting in decreased mitochondrial performance and reduced fitness. This hybrid breakdown is an important component of outbreeding depression and early-stage reproductive isolation.

However, the mechanisms contributing to mitonuclear interactions remain poorly resolved. Here we scored variation in developmental rate (a proxy for fitness) among reciprocal F_2_ inter-population hybrids of the intertidal copepod *Tigriopus californicus*, and used RNA sequencing to assess differences in gene expression between fast- and slow-developing hybrids. In total, differences in expression associated with developmental rate were detected for 2,925 genes, whereas only 135 genes were differentially expressed as a result of differences in mitochondrial genotype. Up-regulated expression in fast developers was enriched for genes involved in chitin-based cuticle development, oxidation-reduction processes, hydrogen peroxide catabolic processes and mitochondrial respiratory chain complex I. In contrast, up-regulation in slow developers was enriched for DNA replication, cell division, DNA damage and DNA repair. Eighty-four nuclear-encoded mitochondrial genes were differentially expressed between fast- and slow-developing copepods, including twelve subunits of the electron transport system (ETS) which all had higher expression in fast developers than in slow developers. Nine of these genes were subunits of ETS complex I. Our results emphasize the major roles that mitonuclear interactions within the ETS, particularly in complex I, play in hybrid breakdown, and resolve strong candidate genes for involvement in mitonuclear interactions.

## Introduction

Coevolved genetic interactions within a population or species can be disrupted by hybridization, and these disruptions may produce incompatibilities that cause loss of fitness in hybrids (i.e., hybrid breakdown; e.g., Burton, 1990; Burton et al., 2006; Ellison & Burton, 2008b; Meiklejohn et al., 2013), potentially contributing to outbreeding depression, early-stage reproductive isolation and speciation (Gershoni et al., 2009; Burton & Barreto, 2012; Hill, 2016, 2019; Sloan et al., 2017). In eukaryotic organisms, incompatibilities underlying these important evolutionary processes may disproportionately involve interactions between genes encoded in the mitochondrial genome and genes encoded in the nuclear genome (Burton & Barreto, 2012). Two major factors contribute to this potential bias for involvement of mitonuclear incompatibilities. First, rates of evolution are higher for mitochondrial DNA than for nuclear DNA (Lynch, 1997; Wallace, 2010), which increases divergence between the mitochondrial genomes of independent taxa at early stages of isolation; these differences in mitochondrial DNA give rise to intrinsic selection on nuclear-encoded genes to maintain compatible interactions (Burton & Barreto, 2012; Osada & Akashi, 2012; Barreto et al., 2018), resulting in taxon-specific mitonuclear coevolution. Second, mitochondrial functions are intimately reliant on mitonuclear interactions, and since mitochondria play critical roles in eukaryotic cells, including producing of the majority of cellular energy (i.e., ATP), mitochondrial dysfunction as a result of these incompatibilities is often associated with negative fitness consequences (Rand et al., 2004; Lane, 2005; Wallace, 2010; Hill, 2015; Hill et al., 2019).

Despite the potential impact of mitonuclear incompatibilities on hybrid breakdown, the physiological and genetic mechanisms underlying these effects are poorly resolved. Although the mitochondrial genome is small (typically ∼16 kb encoding only 37 genes in most metazoans; e.g., Levin et al., 2014), approximately 1,500 nuclear-encoded genes function within mitochondria (Bar-Yaacov et al., 2012; Hill, 2014, 2017), and at least 180 of these genes closely interact with mitochondrial-encoded proteins or RNAs (Burton & Barreto, 2012; Burton et al., 2013; Hill, 2017). The best characterized examples of mitonuclear incompatibilities occur between protein subunits of the electron transport system (ETS; e.g., Ellison & Burton, 2006; Blier et al., 2001; Pichaud et al., 2019), or between a mitochondrial tRNA and its corresponding nuclear-encoded aminoacyl-tRNA synthetase (Meiklejohn et al., 2013). However, genetic incompatibilities in general can also result in changes in gene expression (Haerty & Singh, 2006; Landry et al., 2007) either directly through incompatible regulatory interactions or indirectly through physiological impacts of incompatibilities that alter the regulation of gene expression (Wittkopp et al., 2004; Graze et al., 2009; McManus et al., 2010; Barreto et al., 2015). Both of these possibilities are relevant to mitonuclear interactions, as nuclear-encoded polymerase complexes are responsible for mitochondrial DNA replication and RNA transcription (e.g., Ellison & Burton, 2008a), and variation in mitochondrial functions create regulatory signals that influence nuclear transcription as a part of ‘crosstalk’ between the genomes (Poyton & McEwen, 1996; Cannino et al., 2007; Horan et al., 2013). Therefore, assessing transcriptome-wide changes in gene expression associated with mitonuclear incompatibilities and hybrid breakdown is a promising avenue to resolve not only pathways underlying the effects of incompatibilities, but also specific genes potentially involved in these interactions.

The intertidal copepod *Tigriopus californicus* is an ideal species to study the effects of mitonuclear incompatibilities. *T. californicus* are found in splashpools along the west coast of North America from Baja California, Mexico to Alaska, USA. Populations are restricted to specific rocky outcrops along the coast (Burton, 1997), which results in substantial mitochondrial and nuclear sequence divergence among populations (Burton & Lee, 1994; Burton 1997; Edmands, 2001; Peterson et al., 2013; Pereira et al., 2016; Barreto et al., 2018). Despite these high levels of divergence, inter-population crosses generate viable hybrid offspring in the laboratory (e.g., Burton, 1986), and signatures of inter-genomic coevolution have been detected for nuclear-encoded mitochondrial (N-mt) genes across several geographically isolated populations (Barreto et al., 2018). Specifically, effects of mitonuclear incompatibilities on oxidative phosphorylation (Ellison & Burton, 2006, 2008b; Healy & Burton, 2020; Han & Barreto, 2021), mitochondrial transcription (Ellison & Burton, 2008a), and the evolution of mitochondrial ribosomal proteins (Barreto & Burton, 2012) have been observed in *T. californicus* hybrids. Recent studies have demonstrated strong effects of mitonuclear incompatibilities by comparing nuclear-allele frequencies between reciprocal F_2_ hybrids with fast or slow developmental rate (a proxy for fitness in *T. californicus*; Burton, 1990), and have identified chromosomes that likely contain loci responsible for these effects (Healy & Burton, 2020; Han & Barreto, 2021). However, the genes underlying these effects, and the relative influences of effects on different cellular pathways remain unknown.

In the current study, we examine physiological and genetic mechanisms underlying mitonuclear incompatibilities by using RNA sequencing (RNA-seq) to compare transcriptome-wide variation in gene expression between fast- and slow-developing F_2_ *T. californicus* hybrids. Our goals were: (1) to assess the extent of variation in gene expression associated with differences in developmental rate, (2) to identify genes that were differentially expressed as a result of variation in mitochondrial genotype, (3) to determine biochemical pathways enriched for these differences in gene expression, and (4) to examine patterns of differential expression for both N-mt genes and mitochondrial-encoded genes.

## Materials and Methods

### Copepod collection, culturing and crossing

*T. californicus* adults were collected from supralittoral tidepools at San Diego, California (SD; 32° 45′ N, 117° 15′ W) and Santa Cruz, California (SC; 36° 56′ N, 122° 2′ W) in the summer of 2019. Large plastic pipettes were used to transfer copepods and tidepool water to 1 L plastic bottles. Bottles were transported to Scripps Institution of Oceanography, University of California San Diego within 24 h of collection, and population-specific laboratory cultures were initiated by dividing the collections into 400 mL glass beakers (250 mL per beaker). Cultures were maintained using filtered seawater (35 psu), and were held in incubators at 20 °C under a 12 h:12 h light:dark photoperiod. Powdered spirulina and live *Tetraselmis chuii* algal cultures were added to the cultures as food once per week, but copepods also consumed natural algal growth within their beakers. Laboratory cultures were maintained under these constant conditions for at least 12 months (∼1 month per generation; e.g., Pereira et al., 2021) prior to the initiation of experimental crosses.

Four experimental hybrid lines were established for each reciprocal cross between the two populations: SD♀ x SC♂(SDxSC; lines A-D) and SC♀ x SD♂(SCxSD; lines E-H). Note that SDxSC and SCxSD lines differ in their mitochondrial genotype, which is generally maternally inherited in *T. californicus* (e.g., Burton et al., 2006, but see Lee & Willett, 2022), whereas population-specific contributions to nuclear genotypes are expected to be equal under a neutral assumption (e.g., Lima & Willett, 2018). Virgin females of each population were obtained by splitting mate-guarding pairs using a fine needle (Burton et al., 1981; Burton, 1985). Lines were started by adding 50 virgin females to 2.5 × 15 cm petri dishes containing ∼200 mL filtered seawater and 50 males of the alternative population. Individuals were allowed to pair and mate haphazardly, and lines were maintained and fed as described above for the laboratory cultures. When gravid females were observed, they were transferred to new dishes (one dish per line), and 28 to 39 gravid P_0_ females were obtained per line. F_1_ offspring hatched naturally into the new dish, and once F_1_ copepodids (juveniles) were visible without magnification, the P_0_ females were removed resulting in an F_1_-only dish for each line. *T. californicus* females produce multiple egg sacs from a single mating, and typically ∼22-32 offspring hatch from each egg sac (e.g., Edmands & Harrison, 2003), meaning each F_1_-only dish contained a minimum of many hundreds of offspring. F_1_ individuals matured and mated haphazardly within their dishes, and the dishes were maintained until one week after gravid F_1_ females were initially observed. At this time, F_2_ developmental trials were started, which avoided inadvertently selecting only the fastest developing F_1_ females as parents in the F_2_ trials and prevented any F_2_ offspring that hatched in the F_1_-only dish from reaching adulthood.

### Developmental rate

Developmental rate measurements for F_2_ offspring from each hybrid line were conducted similarly to those described in Healy & Burton (2020). In brief, mature (red) egg sacs were dissected from 30 haphazardly selected F_1_ females for each line using a fine needle. Egg sacs were transferred individually into wells of 6-well plates containing ∼8 mL filtered seawater. Powdered spirulina was added to the wells, and then the plates were placed in the incubators that were used for the F_1_ crosses (at 20 °C; 12 h:12 h light:dark). F_2_ egg sacs hatched overnight, and offspring development was monitored daily with additional spirulina added every other day. Development in *Tigriopus sp*. involves a distinct metamorphosis between the last naupliar (N6) stage and the first copepodid (C1) stage (Raisuddin et al., 2017), which can be used to score developmental rate as time to metamorphosis. As copepodids appeared in the experimental wells, days post hatch (dph) to metamorphosis was scored for each individual, and copepodids were grouped by dph to metamorphosis in line-specific 2 × 10 cm petri dishes containing ∼50 mL filtered seawater. Developmental rate was scored for 467 to 968 F_2_ copepodids per line.

### RNA isolation and RNA-seq

Stage 1 *Tigriopus sp*. copepodids are very small (∼0.35 mm length [Raisuddin et al., 2007]), and obtaining sufficient RNA for standard RNA-seq library preparations from pools of large numbers of individuals at this stage is impractical. Thus, we allowed our scored F_2_ hybrids to progress approximately two additional copepodid stages through development to the C3 stage (∼0.6 mm length) prior to RNA isolation. This progression was tracked by visual monitoring and by time, as under our experimental conditions stage 3 is reached ∼4-5 days after initial metamorphosis to a stage 1 copepodid (Healy et al., *in prep*.). The 100 fastest and 100 slowest developing stage 3 copepodids from each line were snap frozen in liquid nitrogen and stored at -80 °C, and clear separation between fast and slow developers was achieved for every line (minimum of 1 d). Note that monitoring developmental progression for copepodids that are not held individually is imprecise (Tsuboko-Ishii & Burton, 2018), but monitoring at the culture level was necessary given the number of copepodids scored in our study. As a result, it is possible that small numbers of frozen copepodids were at the C2 or C4 stages rather than the targeted C3 stage.

RNA was isolated from the pools of fast and slow developers from each line using TRI Reagent® (Sigma-Aldrich, St. Louis, MO, USA) following the manufacturer’s instructions with modifications as described in Healy et al. (2019). Genomic DNA contamination was removed with an Invitrogen™ TURBO DNA-*free*™ kit (Thermo Fisher Scientific, Waltham, MA, USA) according to the manufacturer’s instructions, and final RNA concentrations were determined with an Invitrogen™ Qubit™ 2.0 Fluorometer and RNA HS assay kit (Thermo Fisher Scientific). RNA samples were submitted to the University of California San Diego Institute for Genomic Medicine Genomics Center for preparation of mRNA stranded libraries for 100 base pair paired-end RNA-seq. The libraries were sequenced on an Illumina NovaSeq 6000 (Illumina Inc., San Diego, CA, USA), and between 18,165,204 and 31,441,512 paired-end reads were obtained for each sample.

### Data analysis and statistics

All analyses were conducted in *R* v4.2.0 (R Core Team, 2022) with α = 0.05. Variation in log-transformed developmental rate as a result of variation in mitochondrial genotype was assessed with a linear mixed-effects model using the *lmerTest* package v3.1.3 (Kuznetsova et al., 2017) with mitochondrial genotype as a fixed factor and line as a random factor.

RNA-seq reads were trimmed to remove any potential adapter sequences with *Cutadapt* v3.4 (Martin, 2011), and were then mapped to a hybrid genome for SD and SC *T. californicus* prepared using the *T. californicus* reference genome (SD reference genome GenBank: GCA_007210705.1, Barreto et al., 2018), a SC-specific genome from population re-sequencing (Healy & Burton, 2020), and published sequences for the SD and SC mitochondrial genomes (GenBank: DQ913891.2 and DQ917374.1, respectively; Burton et al., 2007). Note the SD and SC genomes were masked such that any ‘N’ position in one genome was also ‘N’ in the other genome to avoid any potential mapping biases due to incomplete re-sequencing (Healy & Burton, 2020); this masking procedure between pairs of *T. californicus* populations generally affects less than 2% of coding sequences (Barreto et al., 2018). Genomic feature annotations (i.e., gene models) for the hybrid reference were prepared from previously published annotations for the reference and mitochondrial genomes (Barreto et al., 2018). Sequencing reads were aligned to the hybrid genome using *STAR* v2.7.8a (Dobin et al., 2013) allowing reads to map to a up to two locations in the hybrid genome (“--outFilterMultimapNmax 2” option) so that reads mapping to conserved regions between SD and SC would be included in expression estimates. Overall mapping rates were between 90.67% and 92.94% per sample with the majority of reads mapping uniquely (82.30 ± 1.28%, μ ± σ), which is consistent with relatively high levels of sequence divergence across the transcriptome between SD and SC *T. californicus* (Barreto et al., 2011; Barreto et al., 2015). Reads were counted with *featureCounts* (Liao et al., 2014) from the *Subread* package v2.0.3 using fractional counting (“-M” and “--fraction” options; i.e., reads that mapped to two genes in the hybrid genome were counted as 0.5), and reads for homologous SD and SC genes in the hybrid genome were summed to allow comparisons of total expression for each gene among our RNA samples from pooled F_2_ hybrids.

Variation in transcriptome-wide gene expression patterns was assessed with a principal component analysis (PCA) using the prcomp function from the *R* package *stats* v3.6.2, and 95% confidence ellipses for groups of samples (by developmental rate or mitochondrial genotype) were determined with the *FactoMineR* package v2.4 (Lê et al., 2008). Gene-wise differential expression was assessed by fitting negative bionomial models to the count data with the *edgeR* package v3.38.0 (Robinson et al., 2010) such that main effects of developmental rate and mitochondrial genotype, and interactive effects of these main factors could be tested as described in Lin et al. (2016). In brief, counts were normalized for library size using the relative log expression method (Anders & Huber, 2010; “RLE” option in *edgeR*), low expression genes were filtered with the filterByExp function, dispersions were estimated with the estimateGLMRobustDisp function (Zhou et al., 2014), and factor effects were tested by likelihood ratio tests using the glmFit and glmLRT functions. After filtering, the library sizes in *edgeR* ranged from 14,261,581 to 25,846,327 counts, and differential expression was assessed for 13,994 genes with two analytical approaches. First, we analysed the complete dataset for all eight of our hybrid lines. However, because we pooled the 100 fastest or slowest developers in each line for RNA-seq, the ranges of developmental rate for fast or slow developers had the potential to vary among lines. Thus, we compared fast and slow developmental rates among lines using linear mixed-effects models (as described above for effects of mitochondrial genotype), and performed a second differential expression analysis using a subset of the lines that had similar developmental rates across the copepodid pools sampled for RNA-seq (see Results below).

Previously published gene ontology (GO) functional annotations of the gene models in the *T. californicus* genome were obtained from Barreto et al. (2018; 35,947 annotations for 9,362 genes). We expanded these annotations by running the transcriptome through the *Trinotate* v3.2.2 pipeline (e.g., Bryant et al., 2017), and by manually annotating any remaining unannotated mitochondrial-encoded genes using information from other arthropods (*Aedes aegypti* and *Drosophila melanogaster*) available in the UniProt database (www.uniprot.org). After combining annotations from all sources and removing duplicate annotations, our GO database contained 152,010 annotations for 12,206 genes in the *T. californicus* genome. Functional enrichment analyses were conducted for the differentially expressed (DE) genes with the *goseq* package v1.48.0 (Young et al., 2010) in *R*. GO terms with less than 10 annotations in the *T. californicus* genome were removed from the database (leaving 2,644 GO terms with 119,094 annotations for 12,160 genes) to provide robust tests for functional enrichments, and analyses were run for genes up-regulated in fast developers, genes up-regulated in slow developers, genes differentially expressed between mitochondrial genotypes, and genes with a significant interaction between mitochondrial genotype and developmental rate.

False-discovery rate (FDR) corrections were made for all statistical results from the differential expression and functional enrichment analyses using the Benjamini-Hochberg method (Benjamini & Hochberg, 1995).

## Results

### Developmental rate

Across our hybrid lines metamorphosis was observed from 6 to 22 dph, and median time to metamorphosis ranged from 8 to 10 dph among lines. There was no significant effect of mitochondrial genotype on developmental rate overall (*p* = 0.20; Figure 1), and mitochondrial genotype also did not affect the developmental rates of the 100 fastest or 100 slowest developing copepodids (*p* ≤ 0.20), which were each comprised of 10.9% to 21.4% of the total number of F_2_ copepodids per line. However, despite the lack of effects of mitochondrial genotype, the observed ranges and distributions of developmental rate in “fast” or “slow” developers displayed variation among lines. In particular, for six lines the majority of fast developers metamorphosed by 7 dph (83, 100, 100, 60, 61 and 100% for lines C-H, respectively), whereas for two lines only 4% or 3% of fast developers metamorphosed by 7 dph (lines A and B, respectively). Comparing these two groups of lines, developmental rates in lines A and B were significantly different from developmental rates in lines C-H both overall (*p* = 0.013) and in fast developers (*p* = 1.4 × 10^−3^), but not in slow developers (*p* = 0.060; Figure 2).

**Figure 1.**
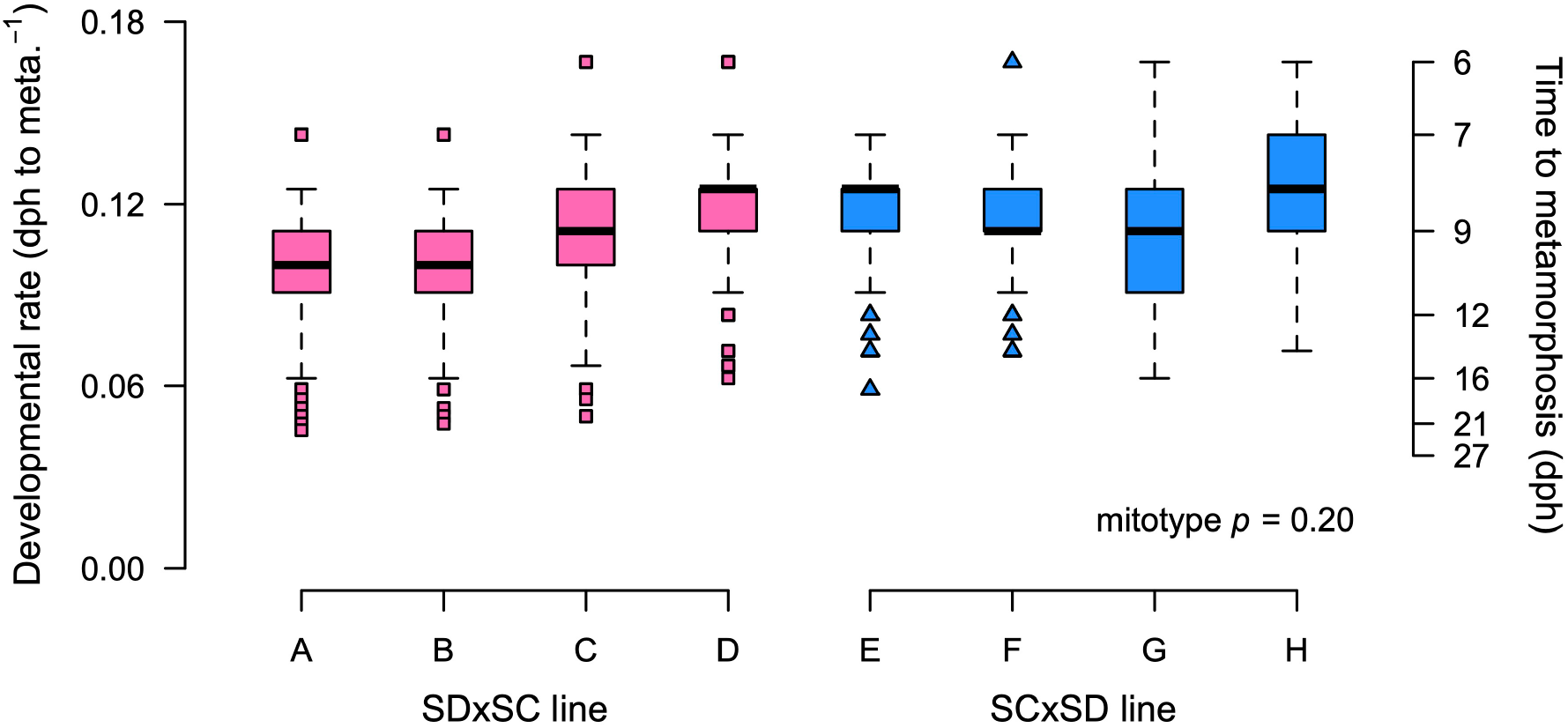
Developmental rate box plots for eight reciprocal F_2_ hybrid lines of *T. californicus* (lines A-D: SDxSC, pink, squares; lines E-H: SCxSD, blue, triangles). Developmental rate is shown both as a rate (left axis) and as the days post hatch (dph) to stage 1 copepodid metamorphosis (right axis).

**Figure 2.**
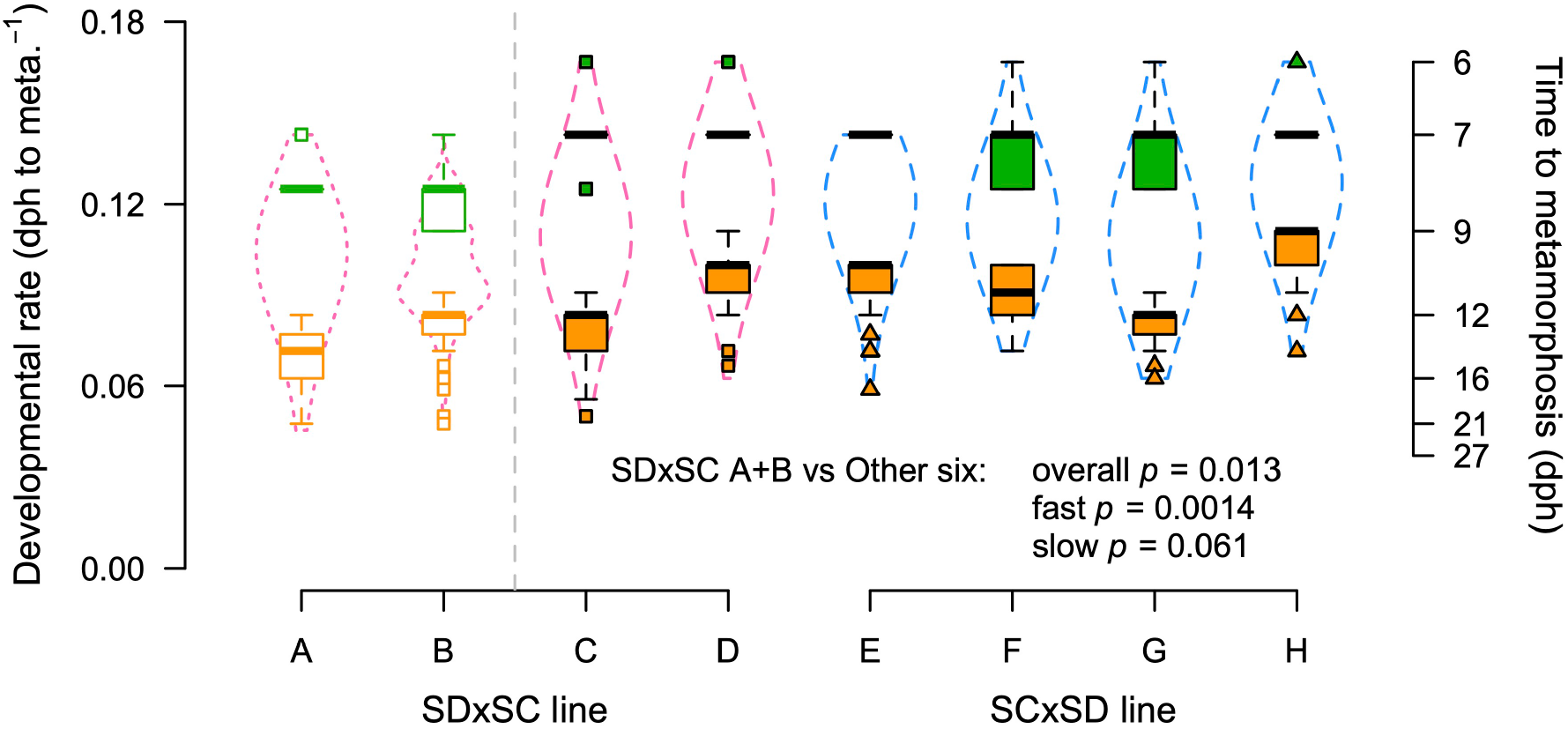
Developmental rate box plots for fast developers (green) and slow developers (orange) from eight reciprocal F_2_ hybrid lines of *T. californicus* (lines A-D: SDxSC, pink density distributions; lines E-H: SCxSD, blue density distributions). Developmental rate is shown both as a rate (left axis) and as the days post hatch (dph) to stage 1 copepodid metamorphosis (right axis). Empty boxes and dotted density distributions show two lines (A and B) with significantly different developmental rates from the other six lines (C-H: filled boxes, dashed density distributions).

Consequently, as discussed above (see Materials and methods), we assessed differential gene expression using all eight lines (hereafter the 8-line DE analysis), and using only lines C-H (hereafter the 6-line DE analysis).

### Transcriptome-wide expression patterns

Gene expression patterns associated with developmental rate were a dominant source of variation across the transcriptome when examined by PCA. The first and second principal components (PC1 and PC2) explained 30.0% and 14.7% of the variation in expression among the RNA-seq samples, respectively, and the fast and slow developers tended to separate along these two axes, particularly along PC1 (slight overlap in 95% confidence ellipses with no overlap at a confidence level of ∼93%; Figure 3a).

**Figure 3.**
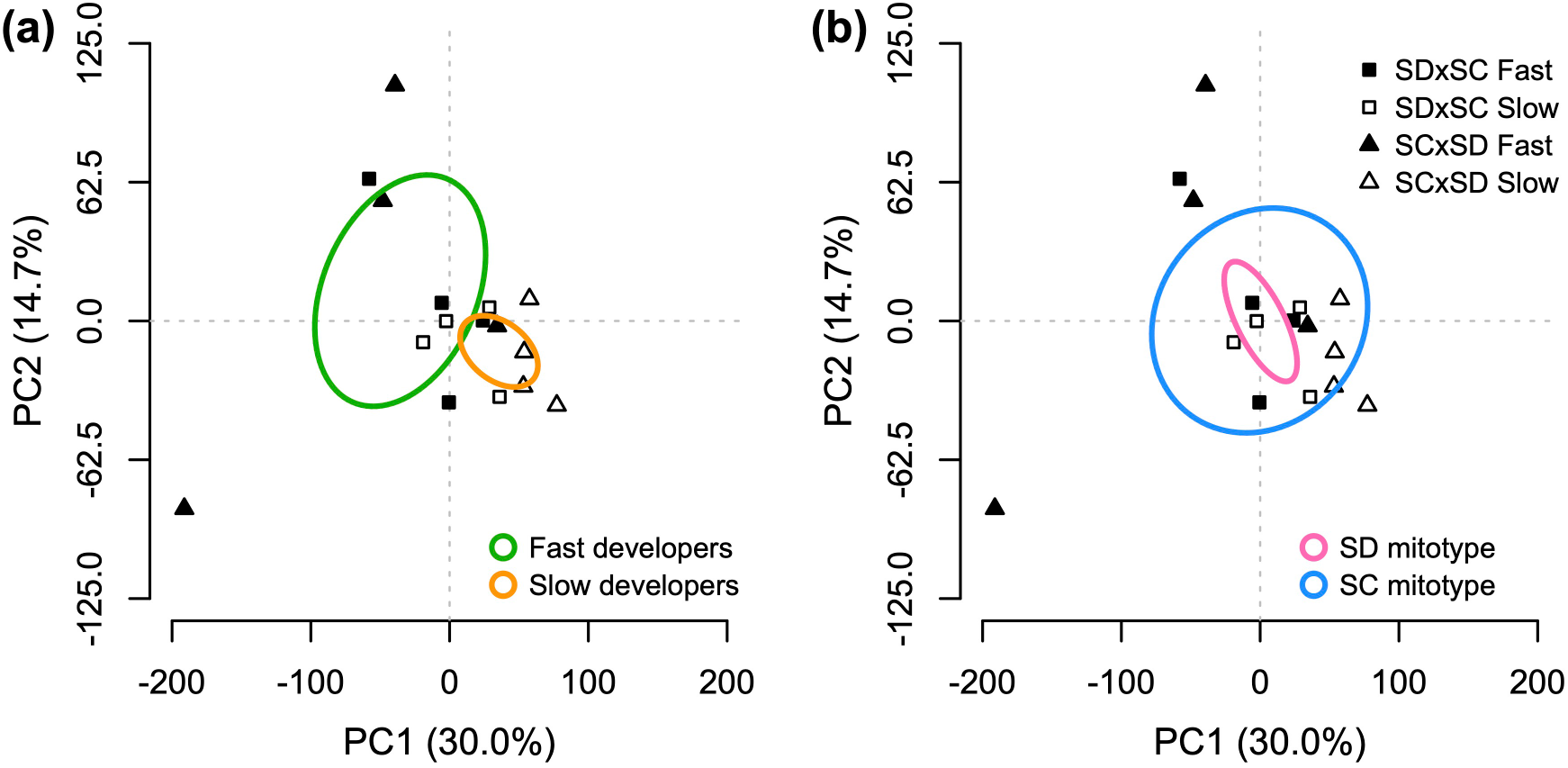
Results from a principal component analysis of transcriptome-wide gene expression for principal components one (PC1) and two (PC2) scores (SDxSC: squares; SCxSD: triangles; fast-developing copepodids: filled symbols; slow-developing copepodids: open symbols). 95% confidence ellipses are shown for developmental rate groups (a – fast developers: green; slow developers: orange) and mitochondrial genotypes (b – SD mitochondrial genotype: pink; SC mitochondrial genotype: blue).

Interestingly, SDxSC lines A and B were the only two lines displaying a positive trajectory from slow to fast developers on PC1, which supports the 6-line DE analysis based on variation in developmental rate among the hybrid lines in our study. Consistent with this pattern, separation between the fast and slow developers along PC1 and PC2 became particularly evident if confidence ellipses were re-calculated for only the 6-line DE analysis RNA-seq samples.

Unlike gene expression associated with developmental rate, little variation associated with mitochondrial genotype was observed along PC1 and PC2 (Figure 3b). Instead, samples from the SDxSC and SCxSD reciprocal lines were separated along PC3 and PC4, which explained 12.7% and 8.4% of the total variation in gene expression, respectively (Figure 4). This indicates relatively modest effects of mitochondrial genotype on transriptome-wide gene expression patterns, particularly in comparison to patterns associated with developmental rate.

**Figure 4.**
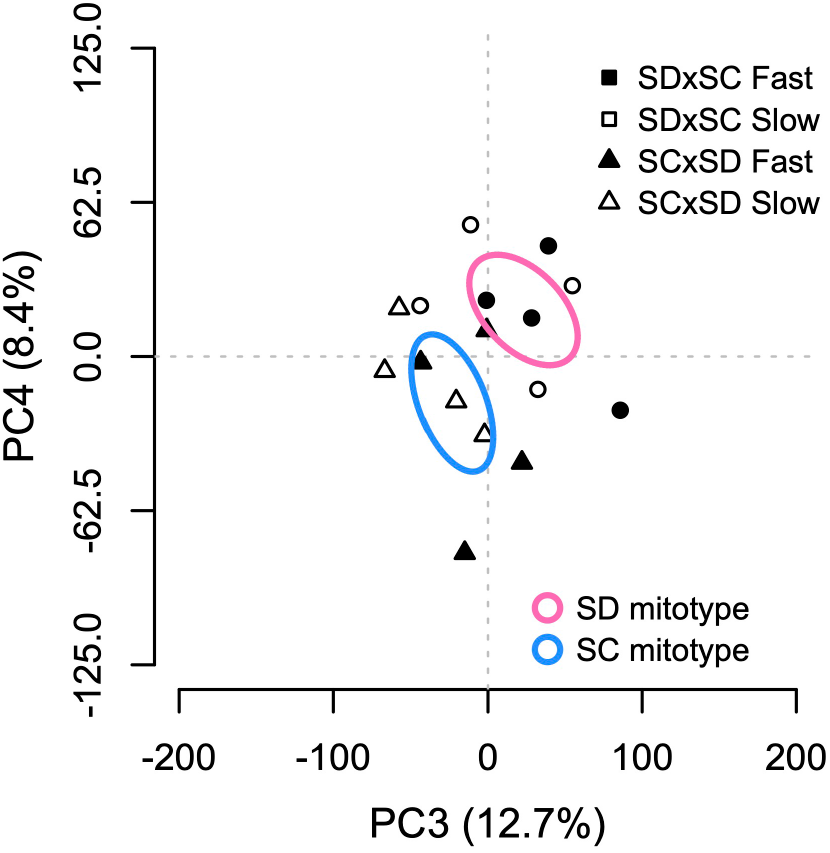
Results from a principal component analysis of transcriptome-wide gene expression for principal components three (PC3) and four (PC4) scores (SDxSC: squares; SCxSD: triangles; fast-developing copepodids: filled symbols; slow-developing copepodids: open symbols). 95% confidence ellipses are shown for mitochondrial genotypes (SD mitochondrial genotype: pink; SC mitochondrial genotype: blue).

### Differential expression associated with developmental rate

Significant differences in gene expression between fast- and slow-developing *T. californicus* were detected for 1,668 genes in the 8-line DE analysis and 2,850 genes in the 6-line DE analysis (Figure 5). In both cases, these relatively high proportions of differential expression across the transcriptome (11.9% or 20.3%) were consistent with the patterns identified by PCA. In general, the 8-line and 6-line DE analyses were highly congruent as 1,593 of the 1,668 DE genes in the 8-line analysis (95.5%) were also detected as DE genes in the 6-line analysis, and there were no strong biases for up- or down-regulation of gene expression in fast or slow developers in either case (900:768 and 1,383:1,467 up-:down-regulated genes in fast developers in the 8-line and 6-line analyses, respectively). However, the distribution of *p*-values versus fold changes in expression was relatively symmetrical around a fold change of zero in the 6-line analysis, whereas this was not the case in the 8-line analysis (Figure 5a,b). In total, 84 N-mt genes were detected as differentially expressed between fast- and slow-developing copepodids (32 common to both the 8-line and 6-line DE analyses, and 1 and 51 unique to each analysis, respectively), whereas no protein-coding genes encoded in the mitochondrial genome demonstrated differential expression associated with developmental rate (FDR-adjusted *p* ≥ 0.36).

**Figure 5.**
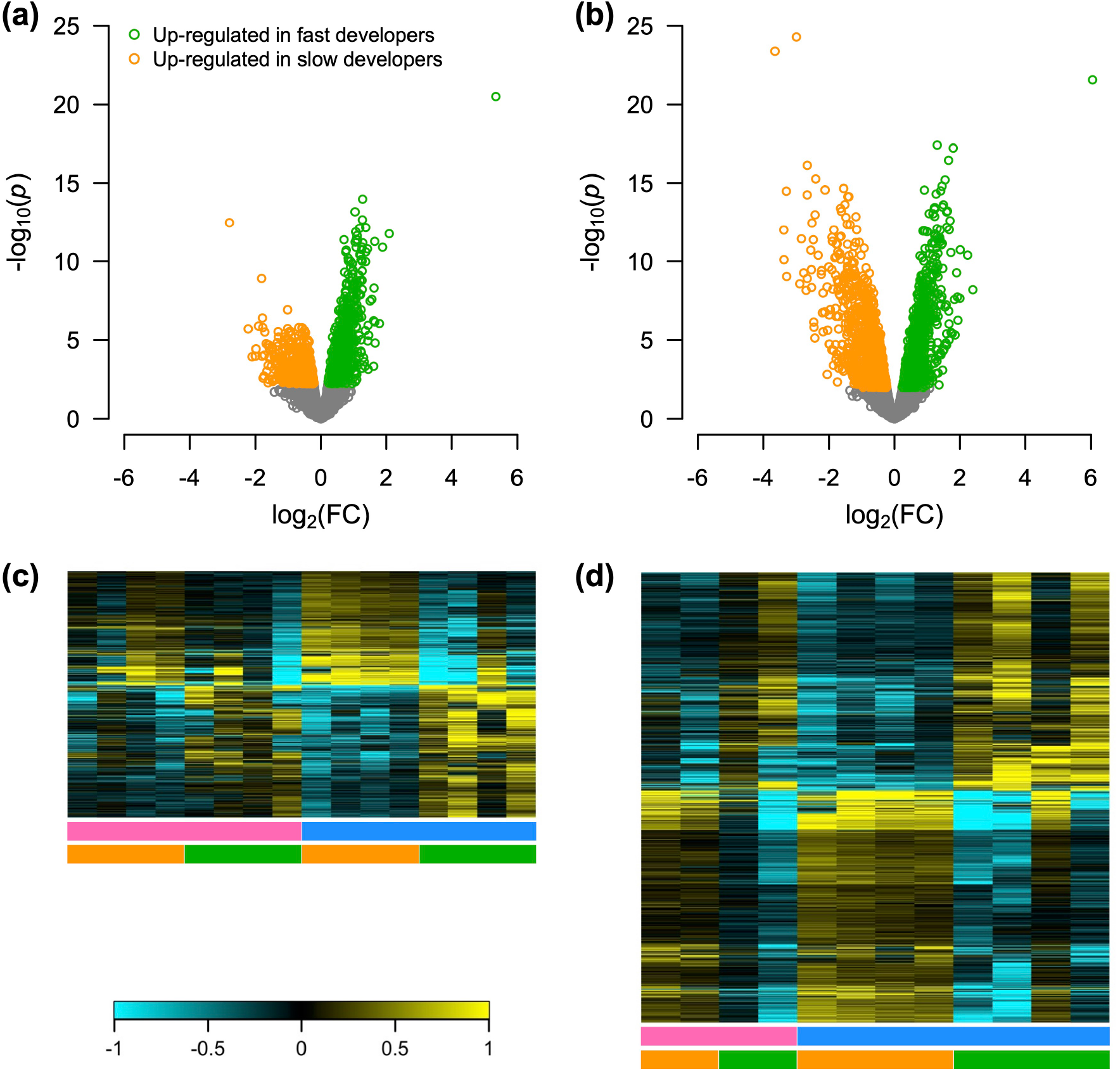
Differences in gene expression between fast- and slow-developing F_2_ hybrid *T. californicus*. Volcano plots (a – 8-line DE analysis; b – 6-line DE analysis) display the negative logarithm of the *p*-value versus the fold change in expression for each gene (green – up-regulated in fast-developing copepodids; orange – up-regulated in slow-developing copepodids; grey – not differentially expressed). Heat maps (c – 8-line DE analysis; d – 6-line DE analysis) display relative variation in expression (higher: yellow; lower: turquoise) among samples (columns) for each differentially expressed gene (rows). Developmental rates and mitochondrial genotypes for each sample are indicated by the horizontal bars below each heat map (developmental rate – fast: green, slow: orange; mitochondrial genotype – SD: pink, SC: blue).

Functional enrichment analyses found 47 GO terms enriched in the genes up-regulated in fast developers (20 common, and 12 and 15 unique from the 8-line and 6-line DE analyses, respectively), and 92 GO terms enriched in the genes up-regulated in slow developers (38 common, and 1 and 53 unique from the 8-line and 6-line DE analyses, respectively). There was variation in the specific GO terms that were enriched among DE genes from the 8-line or 6-line analyses, but the overall patterns regarding cellular functions indicated by the enrichments were similar regardless of which set of DE genes was considered. These results are summarized below using the enrichments for GO biological processes and cellular components.

Enrichments in up-regulated genes were detected for 23 and 46 GO biological process terms in fast- and slow-developing hybrids, respectively (Figure 6). The clearest pattern among these results was the complete lack of overlap in significantly enriched terms between the fast and slow developers. For genes up-regulated in fast-developing copepodids, enrichments were detected related to chitin metabolism (e.g., GO:0040003 and GO:0006030), oxidation-reduction processes (GO:0055114), hydrogen peroxide metabolism (e.g., GO:0042744) and immune responses (e.g., GO:0045087). In contrast, enrichments related to DNA replication (e.g., GO:0006260 and GO:0006270), RNA processing (e.g., GO:0006364 and GO:0006402), cell division (e.g., GO:0051301 and GO:0007049) and DNA repair (e.g., GO:0006281 and GO:0036297) were detected for genes up-regulated in slow-developing copepodids. These major patterns were consistent across the DE genes from the 8-line and 6-line analyses, but additional enriched biological processes for carbohydrate and amino acid metabolism (e.g., GO:0006096 and GO:0006560) were only associated with up-regulation in fast developers using results from the 6-line DE analysis (see details in Figure 6).

**Figure 6.**
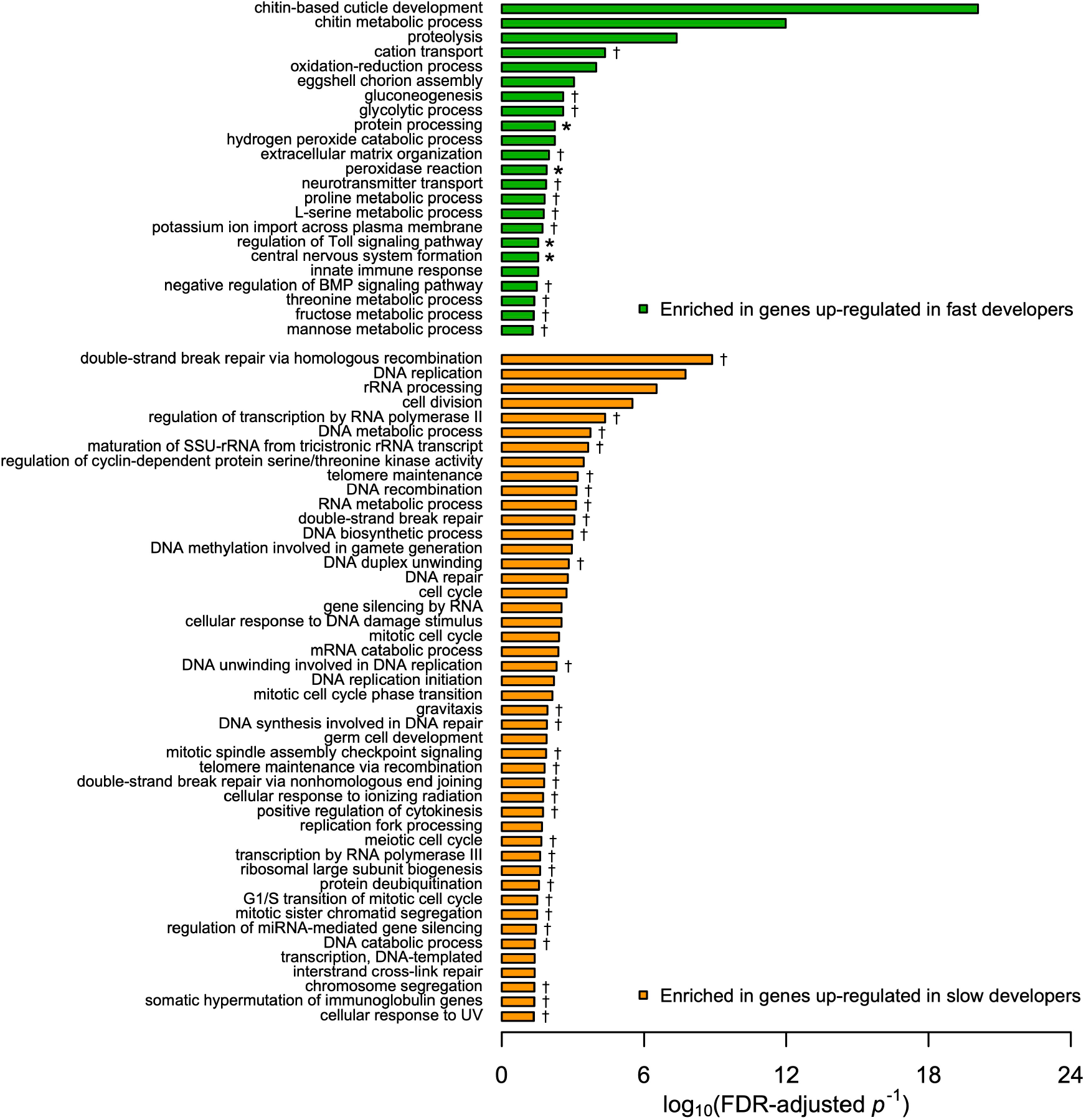
Significantly enriched gene ontology biological process terms among genes up-regulated in fast-developing (green) and slow-developing (orange) copepodids. Bar length indicates the negative logarithm of the FDR-adjusted *p*-value for the enrichment (asterisk – significant for the 8-line DE analysis genes; dagger – significant for the 6-line DE analysis genes; no symbol – significant in genes from both DE analyses; for terms detected in both analyses *p*-values shown are from the 8-line DE analysis).

Compared to biological processes, fewer GO cellular component terms were significantly enriched among DE genes associated with developmental rate. However, the 7 cellular components enriched in genes up-regulated in fast developers, and the 20 cellular components enriched in genes up-regulated in slow developers (Figure 7) suggested similar overall results to those identified based on biological process enrichments. Genes up-regulated in fast developers demonstrated enrichments involving extracellular regions (e.g., GO:0005576 and GO:0005615), whereas genes up-regulated in slow developers were enriched for cellular components related to the nucleus (e.g., GO:0005634 and GO:0005730), a ribosome assembly complex (GO:0032040) and the DNA polymerase complex (GO:0042575). Additionally, among genes significantly up-regulated in fast developers in the 6-line DE analysis, there was an enrichment for the mitochondrial respiratory chain complex I cellular component (GO:0005747). As a result, we specifically examined the DE N-mt genes encoding subunits of the electron transport system (ETS). Twelve nuclear-encoded subunits of the ETS were differentially expressed between fast- and slow-developing hybrid copepodids (FDR-adjusted *p* ≤ 0.048 and fold change in expression 1.21-1.37 for all), and nine of these were subunits of ETS complex I (Table 1).

**Figure 7.**
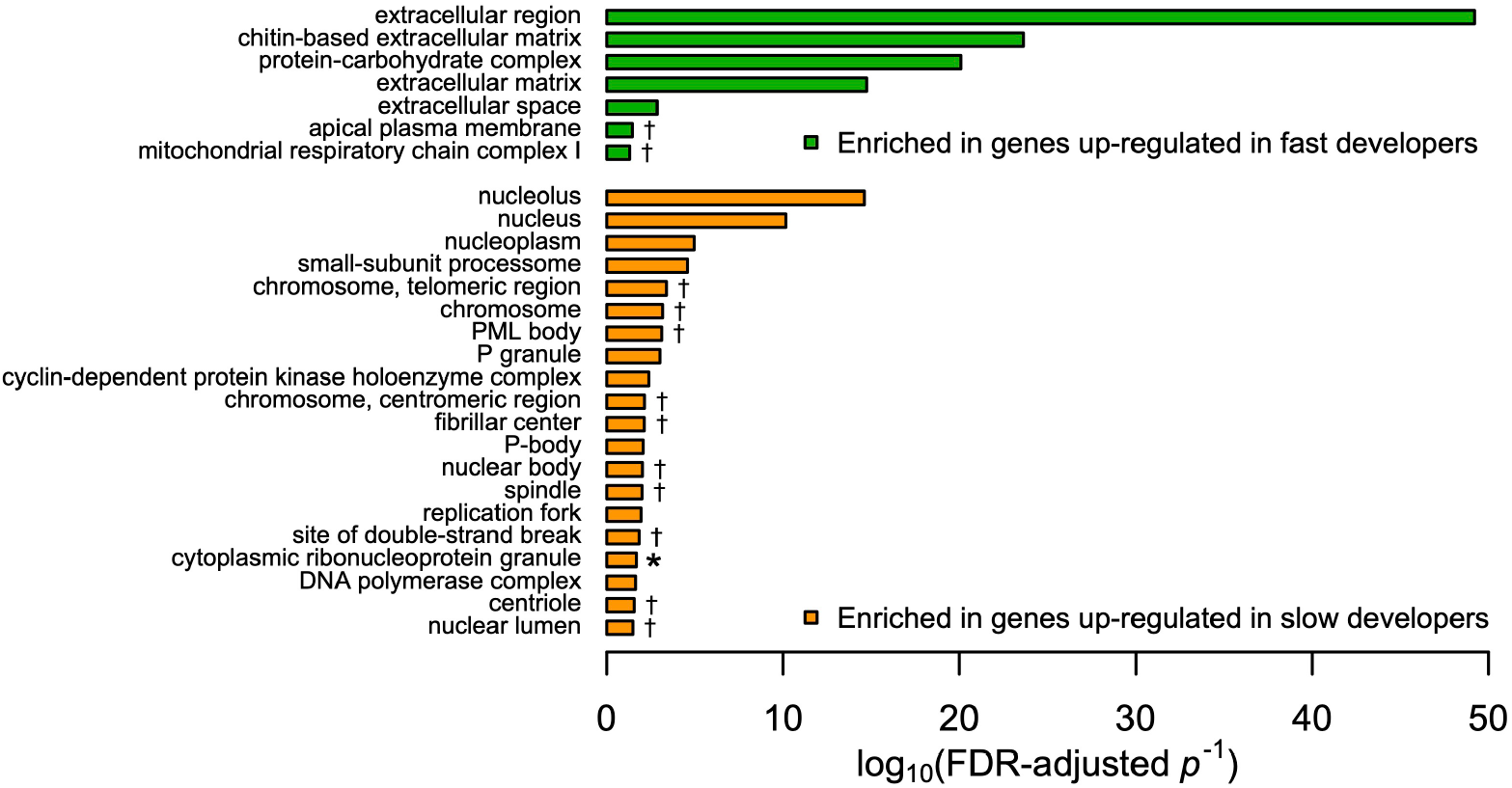
Significantly enriched gene ontology cellular component terms among genes up-regulated in fast-developing (green) and slow-developing (orange) copepodids. Bar length indicates the negative logarithm of the FDR-adjusted *p*-value for the enrichment (asterisk – significant for the 8-line DE analysis genes; dagger – significant for the 6-line DE analysis genes; no symbol – significant in genes from both DE analyses; for terms detected in both analyses *p*-values shown are from the 8-line DE analysis).

**Table 1.** Differences in gene expression between fast- and slow-developing copepodids for the 12 differentially expressed nuclear genes encoding subunits of electron transport system complexes.

| Gene | ETS complex | Chromosome | Counts per million<br>($\mu \pm \text{SEM}$) | | FDR-adjusted $p$ value<sup>†</sup> |
| --- | --- | --- | --- | --- | --- |
|  |  |  | Slow developers | Fast developers |  |
| *ndufs8* | I | 10 | 66.2 ± 1.3 | 81.0 ± 2.9 | 2.0 x 10<sup>-2</sup> |
| *sdhd* | II | 5 | 47.8 ± 1.4 | 59.4 ± 2.7 | 2.1 x 10<sup>-2</sup> |
| *ndufv2* | I | 2 | 77.2 ± 2.0 | 95.8 ± 4.7 | 2.4 x 10<sup>-2</sup> |
| *ndufa10* | I | 6 | 169.3 ± 5.0 | 213.9 ± 10.0 | 2.6 x 10<sup>-2</sup> |
| *atpsyno* | V | 10 | 200.1 ± 6.1 | 257.8 ± 15.0 | 2.6 x 10<sup>-2</sup> |
| *ndufa5* | I | 5 | 41.8 ± 1.1 | 50.4 ± 1.8 | 2.7 x 10<sup>-2</sup> |
| *uqcrq* | III | 3 | 91.0 ± 2.4 | 115.7 ± 7.1 | 3.0 x 10<sup>-2</sup> |
| *ndufs7* | I | 10 | 127.1 ± 2.9 | 159.2 ± 8.7 | 3.3 x 10<sup>-2</sup> |
| *ndufb3* | I | 5 | 53.1 ± 2.1 | 64.8 ± 2.7 | 3.4 x 10<sup>-2</sup> |
| *ndufa8* | I | 3 | 79.9 ± 2.3 | 109.3 ± 13.6 | 3.8 x 10<sup>-2</sup> |
| *ndufa6* | I | 1 | 55.3 ± 1.3 | 69.2 ± 4.6 | 4.2 x 10<sup>-2</sup> |
| *ndufb11* | I | 10 | 72.1 ± 2.4 | 87.5 ± 4.0 | 4.8 x 10<sup>-2</sup> |

<sup>†</sup> from the 6-line DE analysis

### Differential expression associated with mitochondrial genotype

As in most metazoans, mitochondrial DNA is generally maternally inherited in *T. californicus*; however, Lee and Willett (2022) recently detected substantial paternal leakage of mitochondrial DNA in hybrids between some pairs of populations. In contrast, there was virtually no evidence of paternal leakage in our hybrid lines, with 99.8 ± 0.6% (μ ± σ) of the read counts matching the maternal genotype across all samples, suggesting mitochondrial genotypes were maternally inherited in our F_2_ hybrids between the SD and SC populations. In the 8-line DE analysis, 135 genes were significantly differentially expressed between hybrids with a SD mitochondrial genotype (SDxSC) and hybrids with a SC mitochondrial genotype (SCxSD; Figure 8a,b), and 21 genes demonstrated a significant interaction between mitochondrial genotype and developmental rate (Figure 8c). Differences in gene expression as a result of variation in mitochondrial genotype were not evaluated in the 6-line DE analysis, because there was a low sample size for SDxSC lines (2 lines) and an unbalanced design with respect to mitochondrial genotype (2 SDxSC lines [C and D] and 4 SCxSD lines [E-H]).

**Figure 8.**
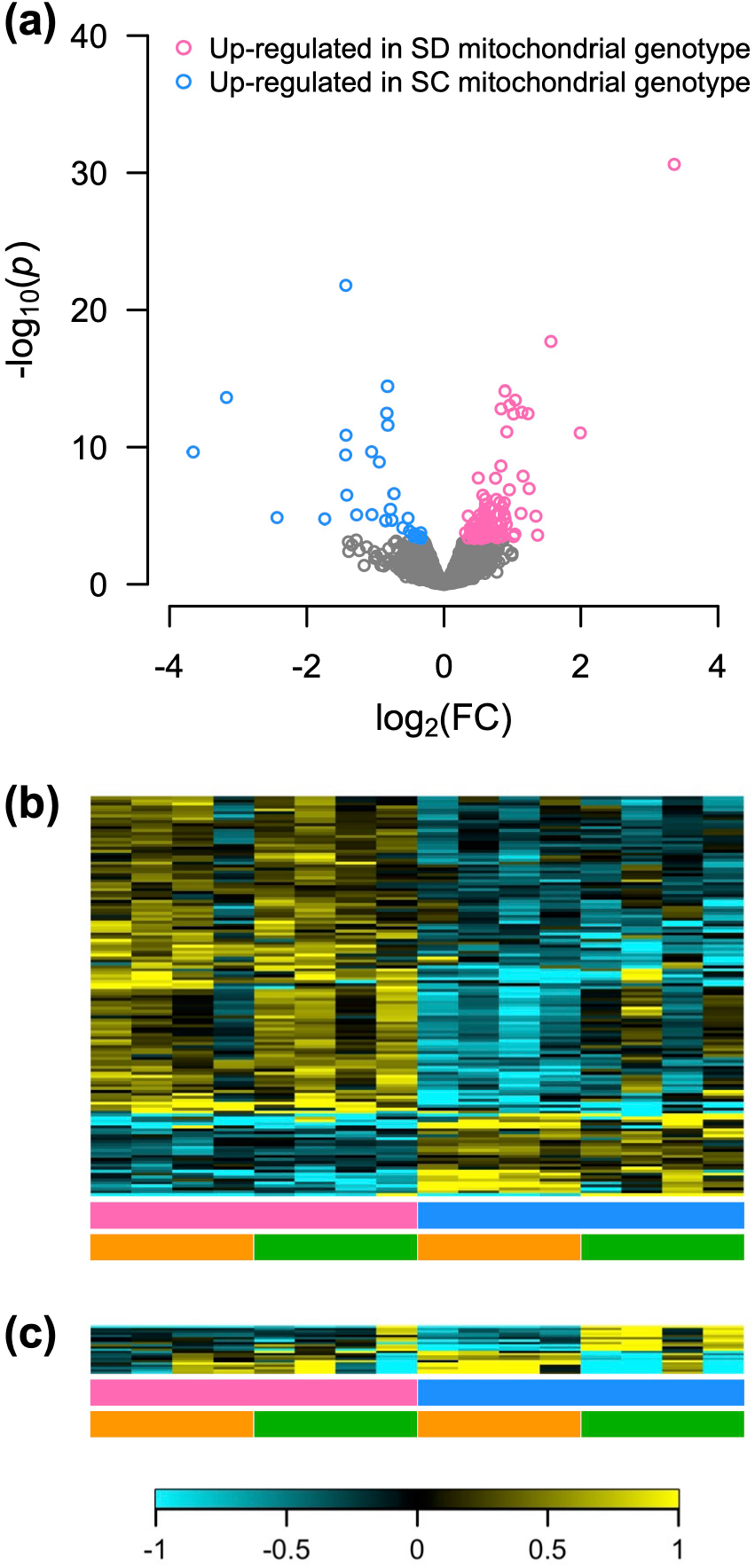
Mitochondrial genotype and interaction effects on gene expression in F_2_ hybrid *T. californicus*. The volcano plot (a) displays the negative logarithm of the *p*-value versus the fold change in expression for each gene differentially expressed between hybrids with SD or SC mitochondrial genotypes (pink – up-regulated in SD mitochondrial genotype; blue – up-regulated SD mitochondrial genotype; grey – not differentially expressed). Heat maps display relative variation in expression (higher: yellow; lower: turquoise) among samples (columns) for each gene (rows) with expression patterns affected by mitochondrial genotype (b) or by a mitochondrial genotype x developmental rate interaction (c). Mitochondrial genotypes and developmental rates for each sample are indicated by the horizontal bars below each heat map (mitochondrial genotype – SD: pink, SC: blue; developmental rate – fast: green, slow: orange).

There were no significant functional enrichments among the DE genes between the SD and SC mitochondrial genotype hybrids, but there was a clear bias in the direction of variation in expression as 107 genes were expressed at higher levels in SDxSC hybrids than in SCxSD hybrids, whereas only 28 genes displayed the opposite pattern (Figure 8a,b). One N-mt gene, 3-hydroxyisobutyrate dehydrogenase (*hibadh*), had higher expression levels in hybrids carrying the SD mitochondrial genotype than in hybrids carrying the SC mitochondrial genotype, and this was the only N-mt gene demonstrating a significant effect of mitochondrial genotype on gene expression. In contrast, 8 protein-coding genes (*mt-nd1, mt-nd3, mt-nd4, mt-nd6, mt-cyb, mt-co1, mt-co3* and *mt-atp8*) and 2 rRNA genes (*mt-rnr1* and *mt-rnr2*) encoded in the mitochondrial genome were differentially expressed between SDxSC and SCxSD hybrids (Table 2). Directions of expression differences among these mitochondrial-encoded genes did not reflect the overall bias of up-regulation of genes in SDxSC hybrids, as 6 DE genes were expressed at higher levels in hybrids with the SC mitochondrial genotype than in hybrids with the SD mitochondrial genotype and 4 DE genes displayed the opposite pattern. The mitochondrial-encoded protein-coding genes all produce subunits of the ETS complexes, but differences in up- or down-regulation for these genes between the two mitochondrial genotypes also did not group consistently based on ETS complex membership (see Table 2).

**Table 2.** Expression levels of mitochondrial-encoded protein- and rRNA-coding genes in SDxSC and SCxSD F_2_ hybrid *T. californicus*.

| Gene<sup>†</sup> | Reads per kilobase per million reads ($\mu \pm \text{SEM}$) | | | | Mitotype<br>FDR-adjusted<br>*p* value |
| --- | --- | --- | --- | --- | --- |
|  | SDxSC slow<br>developers | SDxSC fast<br>developers | SCxSD slow<br>developers | SCxSD fast<br>developers |  |
| *mt-nd1* | 4567 ± 308 | 5001 ± 109 | 2680 ± 124 | 2690 ± 256 | 2.5 x 10<sup>-10</sup> |
| *mt-nd2* | 5483 ± 378 | 5604 ± 204 | 5725 ± 210 | 5699 ± 362 | 0.93 |
| *mt-nd3* | 2068 ± 157 | 2180 ± 93 | 2958 ± 150 | 2980 ± 444 | 3.7 x 10<sup>-2</sup> |
| *mt-nd4* | 2235 ± 161 | 2336 ± 18 | 4356 ± 325 | 3698 ± 276 | 2.5 x 10<sup>-9</sup> |
| *mt-nd4l* | 3763 ± 338 | 4045 ± 116 | 4818 ± 203 | 4565 ± 411 | 0.29 |
| *mt-nd5* | 2621 ± 169 | 2773 ± 44 | 3451 ± 169 | 3127 ± 288 | 0.22 |
| *mt-nd6* | 3913 ± 298 | 4248 ± 131 | 5769 ± 308 | 5122 ± 457 | 3.0 x 10<sup>-2</sup> |
| *mt-cyb* | 3798 ± 236 | 3777 ± 112 | 6519 ± 280 | 6888 ± 406 | 1.3 x 10<sup>-11</sup> |
| *mt-co1* | 8320 ± 445 | 8792 ± 249 | 6071 ± 199 | 6669 ± 542 | 8.5 x 10<sup>-3</sup> |
| *mt-co2* | 6670 ± 468 | 6787 ± 286 | 7493 ± 422 | 7587 ± 505 | 0.64 |
| *mt-co3* | 4098 ± 355 | 4507 ± 238 | 8113 ± 331 | 7370 ± 437 | 4.1 x 10<sup>-10</sup> |
| *mt-atp6* | 10245 ± 773 | 10884 ± 503 | 11840 ± 865 | 11874 ± 1410 | 0.70 |
| *mt-atp8* | 65 ± 9 | 92 ± 14 | 247 ± 33 | 152 ± 28 | 1.1 x 10<sup>-8</sup> |
| *mt-rnr1* | 219 ± 21 | 240 ± 18 | 105 ± 11 | 219 ± 58 | 4.6 x 10<sup>-4</sup> |
| *mt-rnr2* | 206370 ±<br>13569 | 223208 ±<br>5510 | 109165 ±<br>4133 | 130732 ±<br>17155 | 2.3 x 10<sup>-11</sup> |

<sup>†</sup> *nd* – complex I subunit, *cyb* – complex III subunit, *co* – complex IV subunit, *atp* – complex V subunit, and *rnr* – rRNA

In general, genes with expression profiles that were significantly affected by an interaction between mitochondrial genotype and developmental rate demonstrated a consistent pattern when comparing SDxSC and SCxSD hybrids. For all 21 genes with significant interaction effects (Figure 8c), the fold change in expression associated with a difference in developmental rate was higher in hybrids with the SC mitochondrial genotype (7.6 ± 3.5X, mean ± SEM) than in hybrids with the SD mitochondrial genotype (2.7 ± 1.1X), but whether a gene was up- or down-regulated between fast and slow developers was the same regardless of mitochondrial genotype. Genes displaying interactive effects of mitochondrial genotype and developmental rate were significantly enriched for three GO terms: lipid transporter activity (GO:0005319; FDR-adjusted *p* = 2.8 × 10^−5^), extracellular region (GO:0005576; FDR-adjusted *p* = 0.014) and lipid transport (GO:0006869; FDR-adjusted *p* = 0.032); however, these enrichments should be interpreted with some caution given the low number of genes affected by these interactions. No N-mt genes or mitochondrial-encoded genes had differences in expression consistent with effects of mitochondrial genotype by developmental rate interactions.

## Discussion

The expression of genetic incompatibilities in hybrid organisms can result in changes in gene expression through either direct effects of incompatible regulatory interactions or indirect responses to functional consequences of incompatibilities (e.g., Landry et al., 2007; Barreto et al., 2015). The current study reveals high levels of differential gene expression associated with variation in a fitness-related trait (developmental rate) among inter-population F_2_ *T. californicus* hybrids, and recent work in this species has demonstrated that mitonuclear compatibility has a strong effect on variation in developmental rate among these hybrids (Healy & Burton, 2020; Han & Barreto, 2021). Additionally, the slowest developing *T. californicus* hybrids display extreme developmental rates that are outside the ranges of developmental rates observed for offspring from within-population crosses (Healy & Burton, 2020; Han & Barreto, 2021), and consequently the differences in gene expression presented here provide insight into not only the wide range of mechanisms underlying mitonuclear interactions, but also the physiological consequences of these interactions that result in hybrid breakdown.

### Variation in developmental rate among hybrid T. californicus

The developmental rates of inter-population F_2_ *T. californicus* hybrids typically display hybrid breakdown (Burton, 1990; Ellison & Burton, 2008b; Healy & Burton, 2020; Han & Barreto, 2021), and as in previous studies (Ellison & Burton, 2008b; Healy & Burton, 2020), we found that the reciprocal SDxSC and SCxSD hybrids have, on average, similar developmental rates. Despite this, we detected variation in developmental rate among the 8 hybrid lines in the current study (line A and B versus lines C to H). Since only 30 families contributed to each line, the variation in developmental rate among lines may be the result of genetic effects such as random sampling of recombination events or intra-population allelic variants (e.g., Pereira et al., 2016). Alternatively, environmental effects such as differences in algal growth or offspring density may have contributed to the observed variation among lines.

Environmental effects on developmental rate and mitonuclear interactions are common in hybrid organisms in general (Hoekstra et al., 2013, 2018; Baris et al., 2016; Mossman et al., 2016a, 2017; Drummond et al., 2019; Rand & Mossman, 2020; Rand et al., 2022); however, variation in algal growth among lines was minimized manually in the current study, and effects of density dependence on developmental rate are typically minor under our experimental conditions (Healy et al., *in prep*.). Regardless of the cause of variation among lines, strong effects of mitonuclear incompatibilities within lines of hybrid *T. californicus* are expected to result in high degrees of mitonuclear compatibility in fast-developing (high-fitness) hybrids (Healy & Burton, 2020; Han & Barreto, 2021).

### Gene expression differences between high- and low-fitness hybrids

Although ∼1,500 nuclear gene products are potentially imported into mitochondria, 599 N-mt genes have been annotated in the *T. californicus* genome (Barreto et al., 2018), and 84 of these genes were differentially expressed between fast- and slow-developing F_2_ *T. californicus* hybrids. The majority of these (51 genes) were detected only in the 6-line DE analysis, but this is not surprising, because strong signatures of coevolution are only observed in fast-developing hybrids (Healy & Burton, 2020), and developmental rates for the fast developers in the 6-line analysis (lines C-H) were clearly faster than those from the other two lines (lines A and B; Figure 2). Among N-mt genes, there are four groups that most likely involve interactions between mitochondrial- and nuclear-encoded genes: mitochondrial DNA and RNA polymerases, mitochondrial aminoacyl-tRNA synthetases, mitochondrial ribosomal proteins and ETS complex subunits (Burton & Barreto, 2012; Hill, 2015, 2017; Hill et al., 2019). Of the DE N-mt genes in the current study, the clearest association with these groups was the 12 DE genes encoding subunits of the ETS complexes. All of these subunits were expressed at higher levels in fast developers than in slow developers (Table 2), which is consistent with previous studies positively associating ATP synthesis capacity with high fitness in *T. californicus* hybrids (Ellison & Burton, 2006, 2008b; Healy & Burton, 2020; Han & Barreto, 2021). In addition, six other N-mt genes either directly involved in the tricarboxylic acid (TCA) cycle (*idh3b* and *aco2*) or involved in pathways delivering substrates to the TCA cycle (*acss1, pc, mut* and *T05H10*.*6*) were also all up-regulated in fast developers compared to slow developers.

Changes in the proportions of even a small number of interacting mitochondrial proteins can have substantial functional consequences (e.g., Herrmann et al., 2003; Chae et al., 2013), and consequently our results highlight not only specific ETS subunits potentially involved in incompatibilities, but also the key role that dysfunction at complex I may play in hybrid breakdown. Nine of the 12 DE ETS genes encoded subunits of complex I, and the GO cellular component mitochondrial respiratory chain complex I (GO:0005747) was enriched among genes that were up-regulated in fast developers in the 6-line DE analysis. Compared to the other ETS complexes, complex I has the highest number of nuclear-encoded and mitochondrial-encoded subunits (38 and 7 subunits in mammals, respectively; Zhu et al., 2016), and thus is a particularly likely target for the formation of mitonuclear incompatibilities (e.g., Pichaud et al., 2019; Moran et al., 2021). Furthermore, although negative effects of mitonuclear incompatibilities have been demonstrated for complexes I, III and IV in *T. californicus* hybrids (Edmands & Burton, 1999; Willett & Burton, 2001, 2003; Rawson & Burton, 2002; Harrison & Burton, 2006; Ellison & Burton, 2006, 2008b), signatures of divergent selection (i.e., elevated dN/dS) among populations of *T. californicus* are modestly, but significantly, higher for complex I than the other ETS complexes (Barreto et al., 2018).

Potential effects of mitonuclear interactions on mitochondrial transcription (Ellison & Burton, 2008a) and translation (i.e., mitochondrial ribosomal proteins; Barreto & Burton, 2012) have also been detected in *T. californicus*, but relatively few N-mt genes that were differentially expressed between high- and low-fitness hybrids were involved in these functions in the current study. Two mitochondrial ribosomal proteins were differentially expressed between the fast- and slow-developing copepodids with *mrpl19* up-regulated in slow developers and *mrps18c* up-regulated in fast developers, but note neither of these genes were annotated as N-mt genes in Barreto et al. (2018). One mitochondrial aminoacyl-tRNA synthetase (*kars*) and the mitochondrial poly(A) polymerase, *mtpap*, were expressed at higher levels in slow developers than in fast developers, as were N-mt genes involved in mitochondrial DNA replication (*twnk* and *polg*) and translation regulation (*ptcd1* and *guf1*). Additionally, five N-mt genes involved in protein and RNA import into the mitochondria were also all up-regulated in slow developers (*hsp60, tim14, roe1, timm23* and *pnpt1*). Although changes in mRNA levels for individual genes are not necessarily directly related to functional differences at the protein-level (e.g., Hack, 2004), consistent patterns of expression across genes from similar pathways or genes with similar functions are more likely to be indicative of functional effects in the cell.

Mitonuclear incompatibilities and mitochondrial dysfunctions in hybrids have been linked to increased production of reactive oxygen species (ROS) from the ETS (Du et al., 2017; Pichaud et al., 2019), and hybrid lines of *T. californicus* have elevated levels of oxidative damage compared to within-population lines (Barreto & Burton, 2013). The ETS produces ROS as a byproduct of oxidative metabolism, particularly at complex I and III (Brand, 2010; Andreyev et al., 2015), and although ROS can have important functions in cellular signaling, including ‘crosstalk’ between the mitochondria and nucleus (Yun & Finkel, 2014; Shadel & Horvath, 2015), excessive ROS production leads to oxidative stress that is harmful for macromolecules such as DNA (e.g., Temple et al., 2005). In the current study, GO terms associated with antioxidant defense processes such as hydrogen peroxide catabolic process and peroxidase reaction were enriched among genes up-regulated in fast developers, whereas GO terms associated with DNA damage and repair including, for example, DNA repair, cellular response to DNA damage stimulus and interstrand cross-link repair were enriched among genes up-regulated in slow developers. These differences suggest there may be variation in oxidative stress mitigation and damage between the fast and slow developers, potentially indicating additional consequences of mitonuclear incompatibilities associated with the ETS. Consistent with this possibility, hallmark antioxidant enzymes (e.g., Yoo et al., 2020) such as superoxide dismutase (*sod1*) and glutathione *S*-transferase (*mgst1*) were up-regulated in fast-developing copepodids in the current study, and high-fitness *T. californicus* hybrids tend to have lower oxidative damage and higher mitonuclear compatibility than low-fitness hybrids (Barreto & Burton, 2013; Healy & Burton, 2020).

Other functional enrichments among the DE genes in the current study were less clearly connected to potential effects of mitonuclear incompatibilities. For example, enriched up-regulation of cuticle proteins in fast developers could be plausibly connected to the chitin-based exoskeleton in *T. californicus* and the five moults required to reach adulthood from the C1 stage, but connections between chitin metabolic processes and mitonuclear interactions are not readily apparent, despite transgressive expression levels of these proteins generally in *T. californicus* hybrids (Barreto et al., 2015). Cytosolic ribosomal proteins also display transgressive expression patterns in *T. californicus* (Barreto et al., 2015), but no cytosolic ribosomal proteins were differentially expressed between fast and slow developers in the current study. This is at least somewhat surprisingly given the substantial impacts of rates of protein synthesis and variation in ribosomal protein expression on rapid growth during development in Pacific oyster (*Crassostrea gigas*; Hedgecock et al., 2007; Meyer & Manahan, 2010; Pan et al., 2018), and the key role of ribosome abundance in high growth rates in *Saccharomyces sp*. (Warner, 1999; Regenberg et al., 2006; Airoldi et al., 2009).

Taken together, the genes differentially expressed between fast- and slow-developing F_2_ *T. californicus* hybrids suggest a consistent hypothesis for the mechanisms underlying strong effects of mitonuclear incompatibilities in this species. Oxidative phosphorylation functions efficiently at high rates in high-fitness hybrids, whereas incompatible mitonuclear genotypes in low-fitness hybrids cause ETS dysfunction that is associated with increased signals of oxidative damage and potentially signals of compensatory increases in mitochondrial translation and protein import. Although clearly this is largely speculative based on the results of the current study alone, physiological studies in *T. californicus* generally support various aspects of this hypothesis (e.g., Ellison & Burton, 2006, 2008b; Barreto & Burton, 2013; Healy & Burton, 2020), suggesting that further functional work to test these ideas, particularly in the context of variation in developmental rate, is warranted.

### Effects of mitochondrial genotype on the transcriptome

Relatively few genes were differentially expressed between hybrids carrying a SD or SC mitochondrial genotype, which is unlike the substantial variation in nuclear gene expression associated with mitochondrial substitutions in *Drosophila sp*. (Mossman et al., 2016b, 2017, 2019) or differences in mitochondrial genotype in horseshoe bats (*Rhinolophus affinis*; Ding et al., 2021). However, modest effects of mitochondrial genotype on the transcriptome are observed in Atlantic killifish (*Fundulus heteroclitus*) from natural populations with genetic admixture between two subspecies (Flight et al., 2011; Healy et al., 2017), and only small numbers of genes are differentially expressed between *T. californicus* from the SD and SC populations (Barreto et al., 2015).

Interestingly, mitochondrial-encoded genes also display similar expression levels between the SD and SC populations (differences only for *mt-co1* and *mt-cyb* which were more highly expressed in SC than in SD; Barreto et al., 2015), whereas in the current study the clearest group of DE genes between SDxSC and SCxSD hybrids were 10 mitochondrial-encoded genes. Given this difference, it is possible that nuclear genetic background alters mitochondrial transcription in *T. californicus*, as in other species (e.g., Mossman et al., 2016b), which may include direct effects of incompatible mitonuclear regulatory interactions similar to those detected by Ellison & Burton (2008a).

### Mechanisms underlying mitonuclear interactions

Several studies have investigated genetic mechanisms underlying mitonuclear interactions by assessing variation in nuclear allele frequencies in reciprocal *T. californicus* hybrids (Pritchard et al., 2011; Foley et al., 2013; Lima & Willett, 2018; Lima et al., 2019; Healy & Burton, 2020; Han & Barreto, 2020; Pereira et al., 2021). In general, allele frequency deviations away from neutral expectations (i.e., 0.5) reveal consequences of both nuclear-nuclear and mitonuclear interactions with a bias towards the latter, especially in recent studies that focus on variation between reciprocal high-fitness hybrids (Healy & Burton, 2020; Han & Barreto, 2021; Pereira et al., 2021). In particular, fitness differences among hybrids between SD and SC, scored by developmental rate, highlight strong effects of mitonuclear interactions involving loci on chromosomes 1 to 5 (Healy & Burton, 2020). There are 249 annotated N-mt genes are located on chromosomes 1 to 5 in *T. californicus* (47, 71, 42, 46 and 43 in order for chromosomes 1 to 5), and 42 were differentially expressed between fast and slow developers in the results presented here (14, 5, 4, 9 and 10 in order for chromosomes 1 to 5). Additionally, the main findings of the current study indicate that mitonuclear incompatibilities in the ETS complexes likely play key roles underlying variation in developmental rate among F_2_ hybrids between SD and SC, and 22 N-mt ETS subunits are encoded on chromosomes 1 to 5 of which 7 were differentially expressed between the fast and slow developers. Although they are unlikely to be the only nuclear genes involved in mitonuclear interactions, the 6 of these genes that are not subunits of complex II, which has no mitochondrial-encoded subunits (e.g., Saraste, 1999), are currently the strongest candidate genes to underlie mitonuclear incompatibilities in *T. californicus* (complex I: *ndufa6, ndufv2, ndufa8, ndufa5* and *ndufb3*, and complex III: *uqcrq*; Table 1). The relatively large number of genes encoding subunits of complex I among these results further highlight the potential key impacts of incompatibilities in this ETS complex. Therefore, it is possible that interacting subunits of complex I, including the specific subunits identified here, may be potential candidates underlying mitonuclear interactions in eukaryotic organisms more generally as well.

## Acknowledgements

Funding for the current study was provided by a National Science Foundation grant to RSB (IOS1754347). This publication includes data generated at the UC San Diego IGM Genomics Center utilizing an Illumina NovaSeq 6000 that was purchased with funding from a National Institutes of Health SIG grant (#S10 OD026929).

The authors thank Alexis Cody Hargadon for assistance with copepod maintenance, and Dr. Kevin Olsen for comments on the manuscript.

## Data Availability

Prior to final publication, RNA-seq reads will be uploaded to the National Center for Biotechnology Information Sequence Read Archive, and all datasets, the full RNA-seq and enrichment statistical outputs, our annotated hybrid reference for SD and SC, our *T. californicus* GO database, and the genomic locations of the annotated N-mt genes in *T. californicus* will be uploaded to the Dryad Digital Repository.

## Author Contributions

TMH and RSB conceived and designed the experiments; TMH conducted all experiments and analyses; RSB supervised the study; TMH prepared the figures and wrote the manuscript, and TMH and RSB edited and finalized the manuscript.

